# Ferlin C2A-C2B linkers are alternatively spliced, intrinsically disordered, and interact with negatively charged membranes

**DOI:** 10.1101/2025.03.15.643383

**Authors:** Ethiene Kwok, Patricia Khuu, Patrick Reardon, Juan Vanegas, Colin P. Johnson

## Abstract

Ferlins are vesicle trafficking proteins composed of folded C2 domains conjugated by linkers which are largely disordered. Although a role for the for the C2 domains as calcium sensors has been established it remains unclear whether the linkers function beyond acting as passive spacers. We examined the C2A-C2B linker of vertebrate ferlins and found both putative AP2 and SH3 binding short linear motifs (SLiMs) as well as membrane binding sequences for members of the protein family. Specifically for otoferlin we identified an arginine-rich region proximal to a AP2 binding dileucine motif which interacts with negatively charged lipid membranes. Further, the linker region dominated the liposome binding properties of a larger C2A-C2B two-C2 domain segment of otoferlin, suggesting a dominant role in mediating the membrane binding property of the N-terminus. We also found that alternative splicing of the otoferlin C2A-C2B linker adds and additional membrane binding segment and alters the affinity and kinetics of membrane binding. By contrast alternative splicing of the dysferlin linker is not predicted to alter membrane binding but rather alters the number of predicted short linear motifs (SLiMs). In addition we found the otoferlin linker-membrane interaction was sensitive to ionic strength, and simulations suggest positively charged residues including an arginine-rich region mediates binding. We conclude that the C2A-C2B linker of vertebrate ferlins encode both SLiMs which recruit endocytic proteins as well as membrane binding regions that would place the endocytic binding motif proximal to the membrane surface to facilitate endocytosis and synaptic vesicle resupply.

## Introduction

Intrinsically disordered regions (IDR) are protein sequences which lack a well-defined structure and typically populate an ensemble of conformations.^1^ This enhanced flexibility allows for functions that structured domains cannot easily achieve. For example IDRs can enhance binding interactions by extending to capture a target protein using a fly-casting mechanism.^2^ Alternatively, some vesicle trafficking proteins contain disordered sequences which bind and promote membrane budding.^3^ However for many proteins the contribution of the IDR remains uncharacterized. An example is the sensory hair cell protein otoferlin, which resides on synaptic vesicles and contributes to exocytosis and endocytosis at the ribbon synapse. Otoferlin consists of multiple folded C2 domains (denoted C2A-C2G) bridged by long intrinsically disordered linkers.^4^ While the membrane and Ca2+ binding properties of the folded C2 domains have been established, it is unknown whether the IDRs between domains serve a functional role beyond acting as a passive spacer.^5,6^ Otoferlin is one of six ferlin genes within mammals, and while the presence of the disordered linkers is conserved among family members, the sequences vary. Further, splicing of disordered linker regions within ferlins have been reported, including otoferlin and the muscle-enriched paralogue dysferlin. However although the use of alternative sequences within the linkers suggests functional activity, no study has specifically examined the properties of any ferlin linker. In this study we examine the C2A-C2B linker of vertebrate ferlins using a combination of coarse-grained simulation, recombinant protein assays, and solution NMR.

## Materials and Methods

### Protein Production and Purification

pcDNA3.1 otoferlin plasmid (gift from C. Petit, Institut Pasteur et Université Pierre et Marie Curie, France) and pcDNA4/TO/mGFP-dysferlin-myc-his (gift from K. Bushby Newcastle, U.K) were used as templates for amplification of Mus Musculus otoferlin (GenBank: AY586513.1) and Homo sapiens dysferlin (GenBank: AF075575.1), respectively. The relevant sequences were cloned into a pET28a (+) vector (Novagen) between the BamHI and HindIII restriction sites with a TEV sequence inserted between the linker and polyhistidine sequences. BL21-CodonPlus (Agilent) cells containing the expression plasmid were cultured overnight at 37°C in Luria-Bertani broth containing p1% w/v glucose and 50 µg/mL kanamycin and 25 µg/mL chloramphenicol were used to seed 1 liter cultures of Luria-Bertaini broth with kanamycin. 15N and 15N/13C isotopically labeled protein was produced by bacteria grown in MJ9 media containing 1 g/L 15NH4Cl and 2 g/L of 12C or 13C glucose. Cultures were grown to an optical density of 0.6 at 37°C and induced with 0.5 mM isopropyl β-D1-thiogalactopyranoside (IPTG) for 16 hours at 18°C. Cultures were centrifuged at 4000 rpm at 4°C for 20 minutes and resuspended in lysis buffer: 50 mM HEPES pH 8, 250 mM NaCl, 10% (v/v) glycerol, 5 mM CaCl2, 1 mM phenylmethanesulfonyl fluoride (PMSF), and 1 µM leupeptin, pepstatin A, and aprotinin. Cells were lysed by sonication in four sets of two minutes each. 0.5% CHAPS (w/v) was added to the total lysate and left to rock for 1 hour on ice. Soluble fractions were then obtained by centrifugation in a Beckman J2-21 centrifuge at 20,000 x g at 4°C for 60 minutes. Lysate was bound to HisPur Cobalt resin (Thermo Scientific) for 2 hours with rocking at 4°C. The Cobalt resin was washed with the following buffers at pH 7.5: (a) 50 mM Tris, 1 M NaCl, 5% Glycerol, and (b) 50 mM Tris, 300 mM NaCl, 20 mM imidazole. The bound protein was eluted with buffer containing 50 mM Tris, 300 mM NaCl, and 300 mM imidazole (pH 7.5). Proteins were then purified on a Superdex 75 (GE lifesciences) size exclusion chromatography column with a buffer containing 50 mM Tris and 400 mM NaCl (pH 7.5). Purified proteins were buffer exchanged into 25 mM HEPES and 50 mM NaCl (pH 6.0) using Zeba Spin Desalting Column. Samples were analyzed by Sodium dodecyl sulfate– polyacrylamide gel electrophoresis (SDS–PAGE) gel for purity.

#### Liposome Preparation

Liposomes were prepared as described previously.^6,7^ Briefly, lipids were dissolved in chloroform and dried under vacuum until the solvent was removed. The dried lipids were then rehydrated in a 20mM Tris, 100 mM NaCl pH 7 solution to a concentration of 1 mM and extruded using a membrane with a 50 nm pore size. All materials were purchased from Avanti Polar Lipids.

#### Fluorescence Measurements

Measurements were conducted using a QM-40 (Photon Technology International, Birmingham, NJ). The generalized polarization (GP) value was calculated using GP = (I430 - I480)/(I430 + I480). I430 and I480 are the emission intensities at 430 and 480 nm respectively. Data were collected at a 1.0 nm step size with an integration time of 0.1 s. Each sample was measured multiple times to ensure that the system reached steady state. Fluorescence polarization measurements were conducted using an excitation of 490 nm and emission at 520 nm.

#### NMR Measurements

NMR spectra were collected at 303K on a 800-MHz Bruker Avance IIIHD spectrometer equipped with a TCI cryoprobe. Backbone resonance assignments were determined from a set of [15N, 1H] TROSY based triple-resonance experiments (HNCA, HNCACB, HNCOCACB, HNCO, HNCACO). NMR spectra were processed with NMRPipe and analyzed with CcpNmr Analysis.^8,9^ TALOS-N was used to calculate disorder propensity.^10^ All 3D datasets were collected using non-uniform sampling (NUS), with a fully random sampling pattern. NUS data were reconstructed using SMILE and nmrPipe.^11^

### Sedimentation Assay

Otoferlin or dysferlin protein was mixed with liposomes (100 μg) in buffer (20 mM Tris, pH 7.4, 150 mM NaCl). The mixture was incubated for 10 minutes at 23 °C and centrifuged at 90,000g for 45 min in a TA-100 ultracentrifuge (Beckmann Instruments). Supernatant and pelleted fractions were subsequently analyzed by SDS–PAGE.

### Computational Simulations

Molecular dynamics simulations were performed with the GROMACS package.^12^ All systems were modeled with the coarse-grained SPICA force-field.^13–15^ A custom patch from the SPICA developers (https://github.com/SPICA-group/gromacs-spica) was applied to GROMACS before compilation to enable the SPICA angle potential. The initial structure of the largely disordered linker domains was obtained using AlphaFold 2 (AF2) through a local ColabFold installation (version 1.5.5) which queries the (https://api.colabfold.com) server for the multiple sequence alignment. ^16–19^ The initial structure for the otoferlin C2A domain was obtained from the protein databank (PDBID: 3L9B). All-atom membrane patches with a composition of 75 % POPC and 25 % POPS were first built with the CHARMM-GUI webserver (https://www.charmm-gui.org/).^20,21^ Each membrane leaflet contained 220 lipids (165 POPC and 55 POPS), which resulted in a patch of approx. 120 x 120 Å to accommodate the extended linkers. A linker or C2A domain was then placed in the vicinity of the membrane patch (within 10 Å), but not directly in contact with the lipids during initial setup. A water layer approximately 100 Å thick was added to the system to allow ample space for the proteins to bind and unbind from the membrane. Sodium and chloride ions were added to neutralize the system and have concentration corresponding to 150 mM NaCl. After the complete system was built, the SPICA-tools (https://github.com/SPICA-group/spica-tools) were used to convert the all-atom system to the CG SPICA model.^13,14^ Secondary and tertiary protein structural elements were maintained using and elastic network model with a force constant of 1.195 kcal/Å2 and a cutoff of 9.0 Å. For dysferlin, the AF2 model predicts a short anti-parallel beta-sheet and loop structure at the N-terminus between residues 1 and 19. For isoform 1 and 2, AF2 model predicts a longer anti-parallel beta-sheet structure with residues 105-111 complementary H-bonding residues 121-127 at the C-terminus, which is preceded by an alpha-helical region between residues 82-100.

Electrostatic interactions were computed using the particle-mesh Ewald method with a real space cutoff 1.5 nm and a Fourier grid spacing of 0.5 nm, while Van der Waals interactions were computed using 12-4 or 9-6 Lennard-Jones potentials with a cutoff of 1.5 nm. Newton’s equations of motion were integrated with a leap-frog algorithm using a 20 fs time step. The temperature of the system was held constant at 37 °C using a velocity-rescaling algorithm with a time constant of 1 ps, and the pressure was held constant at 1 atm using a semi-isotropic stochastic cell-rescaling algorithm using a time constant of 5 ps.^22,23^ Five independent replicas were run with different initial membrane configurations. Each replica was first energy minimized for 1,000 steps using a steepest descent algorithm, followed by a 100 ns membrane equilibration period were the protein was under harmonic position restraints using spring constants of 1,000 kJ/mol/nm2. After equilibration, each replica was simulated without any restraints for 2 microseconds. Positions of all atoms were saved to the trajectory in 100 ps intervals. Trajectory analysis was performed with the MDAnalysis python library and plots were generated with Matplotlib.^24,25^ Graphical representations of the CG systems were created with NGLview.^26^

## Results

### Ferlin C2A-C2B linkers encode putative membrane binding and SLiM sequences

To identify disordered regions within ferlin proteins we analyzed the amino acid sequence of otoferlin, dysferlin, myoferlin, Fer1L5, and Fer1L6 using IPRED3 (Fig. 1).^27^ Among several shorter predicted disordered regions we found two long disordered sequences within otoferlin that include the alternatively spliced C2A-C2B linker and a second sequence between C2D and C2F (Fig. 1). Disorder within the C2A-C2B linker was also found for dysferlin, myoferlin, and to a lesser extent for Fer1L5 (Fig. 1, arrow). Although lacking a C2A domain, Fer1L6 also appears to retain a disordered sequence at the N-terminus.

**Figure 1:**
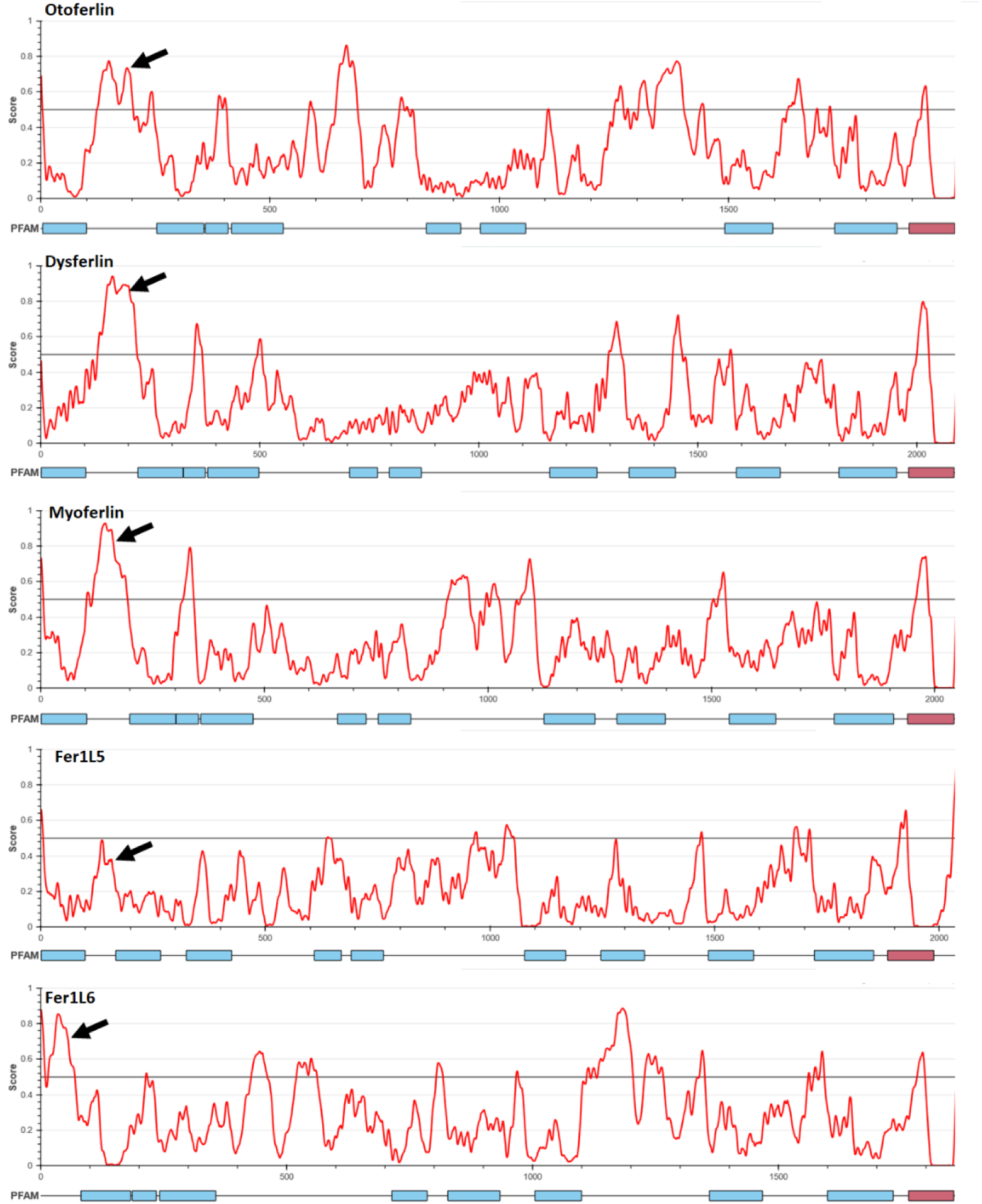
Output of IUPred3 for ferlin proteins otoferlin, dysferlin, myoferlin, Fer1L5, and Fer1L6. Arrow denotes the intrinsically disordered region at the N-terminus

Multiple vertebrate ferlins including otoferlin undergo alternative splicing within the C2A-C2B linker. Examination of the otoferlin cochlear (Q9ESF1, canonical isoform 1) and brain enriched (Q9ESF1-2, isoform 2) linker sequences reveal a disproportionate number of positively charged lysine and arginine residues, and few hydrophobic residues (Fig. 2A). Isoform 2 encodes an additional lysine-rich sequence SKGKEKTKGGRDGEH near the middle of the linker (Fig.2A). Based on the fraction of negative and positively charged residues, both linker sequences reside in the Janus region of a Pappu diagram of states plot, suggesting context dependent conformations (Fig. S1).^28^ Given the enrichment in positively charged residues within the linker isoforms we next assessed the potential of the linker sequences to interact with lipid membranes using Residue-Specific Membrane-Association Propensities of intrinsically disordered proteins (ReSMAP).^29^ Analysis of the linker using ReSMAP revealed a putative membrane binding sequence (a.a. 140-180) located in the middle of the linker region (Fig. 2B). Charged residues compose approximately 35% of the 40 a.a. region identified by ReSMAP, with 20% of the sequence encoding for arginine and lysine. The additional lysine enriched sequence found within isoform 2 generates a second putative membrane binding sequence located proximal to the first binding region (Fig. 2B). The first putative binding region display maxima at residues 168-170 which includes R168 and R169. The second putative binding region within isoform 2 peaks at R187. The identified binding regions appear to be conserved across species, ranging from mouse to zebrafish (Fig. S2). Analysis of the remaining vertebrate ferlin C2A-C2B linkers suggest variability in their membrane binding properties (Fig. S2). For dysferlin (O75923), a small putative binding sequence was found at the extreme C-terminus of the linker proximal to the adjacent C2B domain. Like otoferlin a known dysferlin splice variant adds additional residues within the C2A-C2B linker; however this sequence was not identified by ReSMAP as interacting with membranes. Instead, the inserted residues include a short linear motif (SLiMs) sequence ETWSLL identified using Eukaryotic Linear Motif (ELM) as a putative acidic dileucine AP2 binding motif.^30^ Like dysferlin, alternative splicing of the human myoferlin C2A-C2B linker alters the number of endocytotic SLiMs, with the canonical sequence encoding for up to 6 SH2 and SH3 binding SLiMs, two of which are not found in the alternatively splice linker. We conclude that multiple vertebrate ferlins appear to encode a membrane binding sequence within the C2A-C2B linker, and alternative splicing of otoferlin increases the number of predicted membrane interacting sites while splice variation of the linker in dysferlin alters the number of endocytotic binding motifs within the linker.

**Figure 2:**
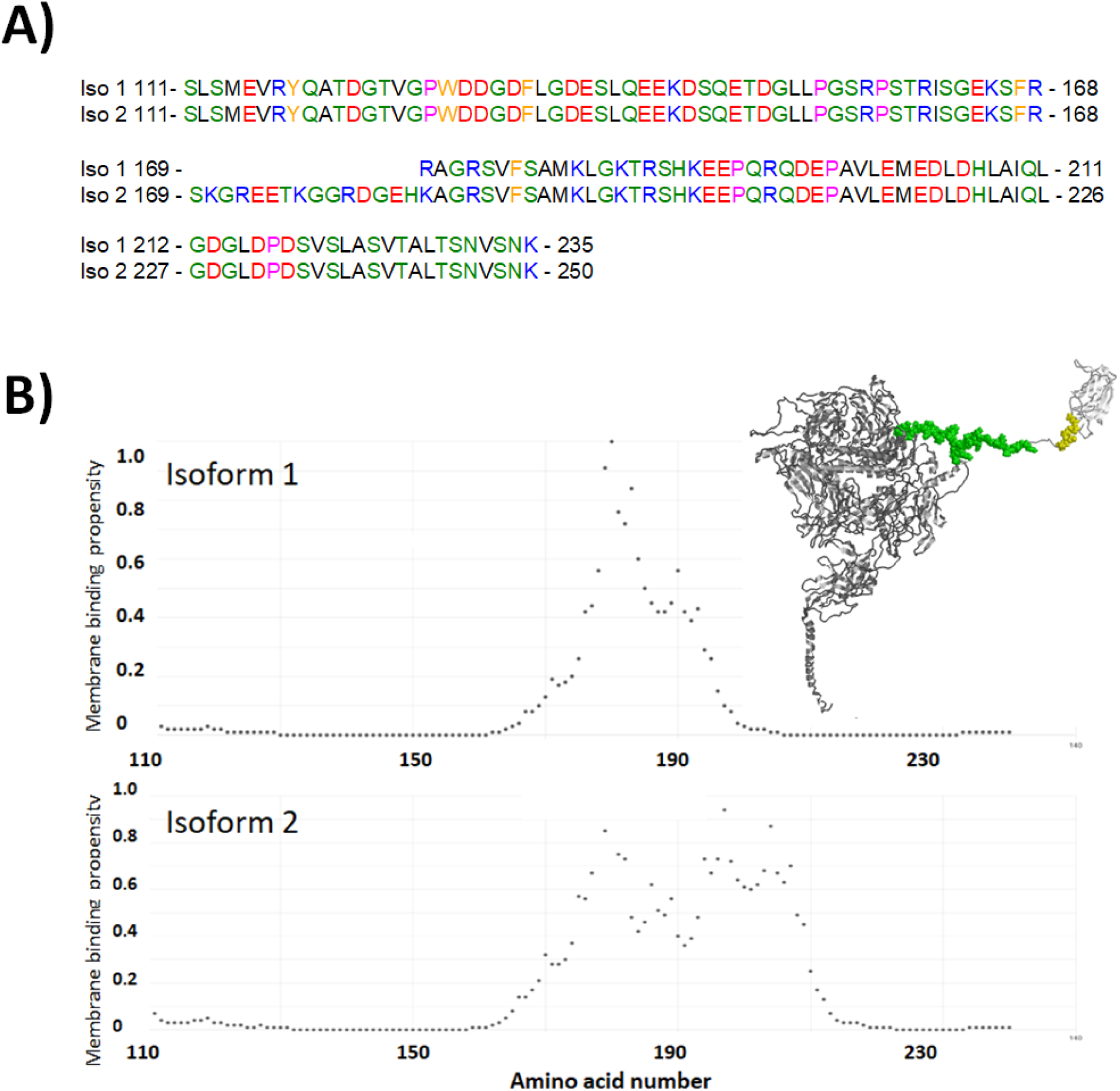
(A) Amino acid sequence alignment of C2A-C2B linker residues for otoferlin Q9ESF1-1 (isoform 1) and Q9ESF1-2 (isoform 2). (B) ReSMAP predicted membrane-binding propensities for otoferlin residues 110-240 for Uniprot Q9ESF1-1 (top panel) and Q9ESF1-2 (bottom panel). Inset displays the Alphfold predicted structure of otoferlin in which the dileucine motif is highlighted in yellow and the predicted membrane binding residues highlighted in green.

### The C2A-C2B linker binds negatively charged membranes

Given the results of the ReSMAP prediction we sought to test whether the C2A-C2B linker binds membranes. We therefore conducted a liposome cosedimentation assay using a recombinant construct composed of otoferlin isoform 2 residues 141-209, encompassing the predicted membrane binding region. When mixed with liposomes composed of 1-palmitoyl-2-oleoyl-sn-glycero-3-phosphocholine (POPC) the recombinant linker did not sediment (Fig. 3A). Given the linker sequence encodes for many charged and polar residues we also tested for linker-membrane interaction using liposomes composed of 20% 1-palmitoyl-2-oleoyl-sn-glycero-3-phospho- L -serine (POPS) and 80% POPC. In contrast to samples containing POPC liposomes, the linker construct cosedimented with negatively charged POPS/POPC liposome samples (Fig 3A,C). We also tested a separate construct composed of the N-terminal C2A domain and proximal linker (amino acids 1-140), which did not cosediment with liposomes regardless of POPS (Fig. 3B,C). In addition, the N-terminal 1-140 construct failed to cosediment when added to liposome samples also containing the membrane binding 141-209 linker region, suggesting the membrane binding region does not recruit the N-terminal 1-140 segment. (Fig. S3A). To determine if the shorter isoform 1 linker also bound membranes we repeated the cosedimentation assay with a recombinant isoform 1 linker and found that like isoform 2, the shorter isoform 1 linker cosedimented with liposomes composed of 20% POPS (Fig. 3C). Qualitatively we also found a recombinant dysferlin C2A-C2B linker composed of residues 101-217 also sedimented with POPS/POPC liposomes albeit to a lesser degree compared to otoferlin (Fig. S3B).

**Figure 3.**
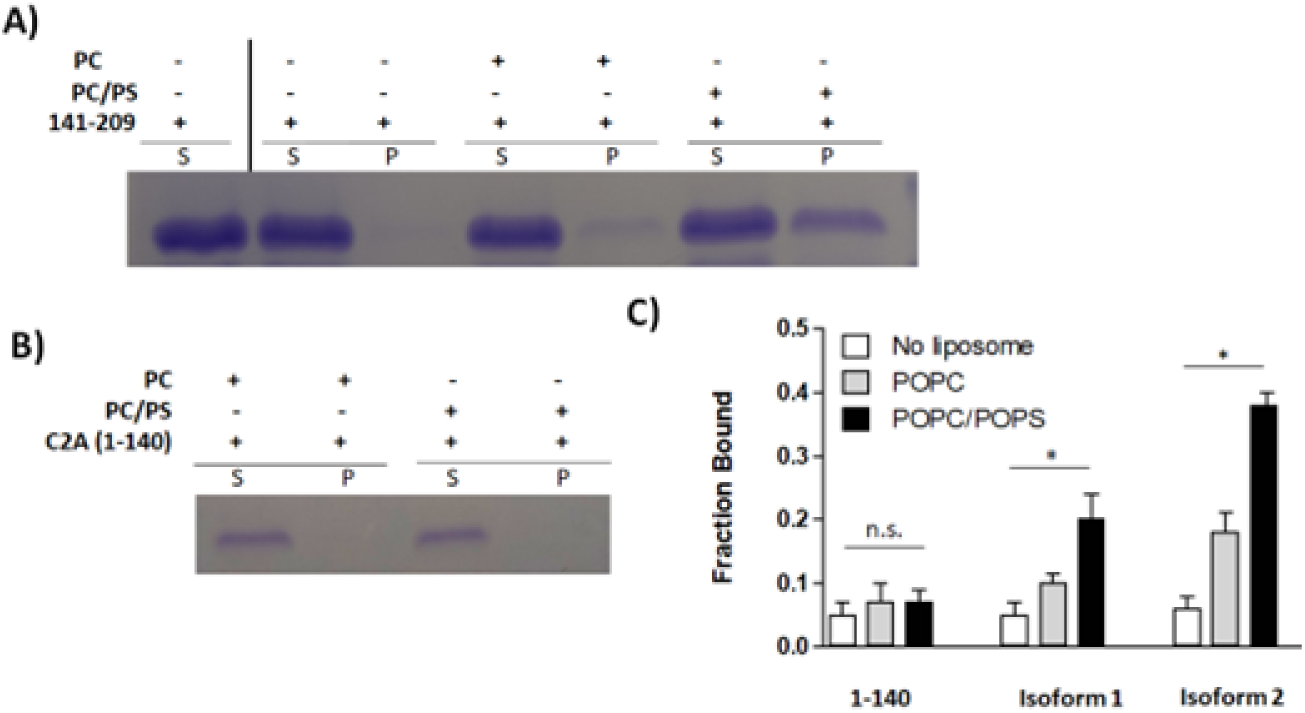
The otoferlin C2A-C2B linker cosediments with liposomes. (A) Representative cosedimentation results for residues 141-209 mixed with POPC liposomes or 20% POPS / 80% POPC liposomes. The first lane represents the protein input, with subsequent lanes for PC and POPC/POPS liposome samples. S denotes the supernatant and P the pellet. (B) The N-terminal residues 1-140 of otoferlin including the C2A domain do not cosediment with liposomes of either PC or PC/PS lipids. (C) Quantitation of the results of the liposome binding assay for 1-140, isoform 1, and isoform 2 linkers. (*t* test, *p* < 0.001). Error bars represent ± standard deviation, *n* = 3.

The ferlin C2A-C2B linkers are predicted to be largely disordered. To experimentally characterize the structure of the linker we collected circular dichroism (CD) measurements of residues of otoferlin isoform 2 as well as dysferlin (Fig. 4). Consistent with a disordered structure we found that the linker spectra exhibited a minimum below 200 nm and was absent of any positive CD signal (Fig. 4A, B). Subsequently we used solution NMR to assign the resonances of C13, N15 isotopically labeled linker residues 141-209 of otoferlin and found the chemical shifts resided in a narrow range of values indicative of a disordered polypeptide, in agreement with the CD spectra (Fig. 4C). Analysis using N-TALOS also indicated the linker is disordered (Fig. S4). As a test for lipid interaction we monitored changes in the chemical shift of N15 labeled 141-209 linker when mixed with POPC/POPS liposomes (Fig.4D). The addition of liposomes to the sample reduced the intensity of resonance peaks for the N15 labeled recombinant linker, consistent with decreased degrees of freedom upon interaction with liposomes.

**Figure 4.**
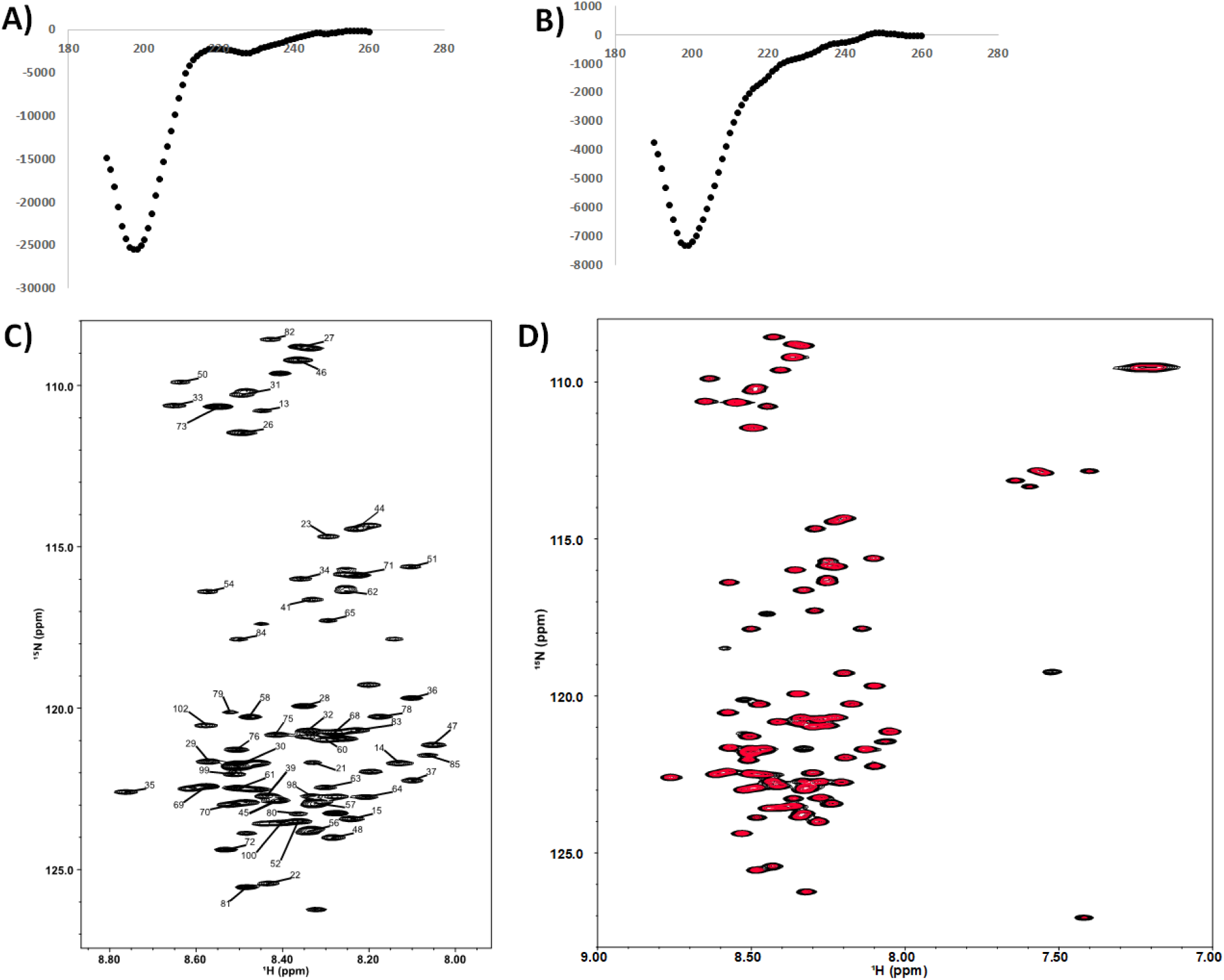
Circular dichroism spectra of otoferlin (A) and (B) dysferlin C2A-C2B linker. (C) 15N -1H spectra of otoferlin residues 141-209. (D) 15N -1H spectra of the linker in the presence of POPS/POPC liposomes. Decreased peak intensity for amino acids influenced by lipids are indicated in red.

As an additional independent method of measuring membrane interaction we monitored binding using laurdan-labeled liposomes. Laurdan is a solvatochromic membrane probe with an emission maximum that shifts upon protein absorption to the liposome surface.^31^ When mixed with liposomes we found the linkers of both otoferlin isoforms shifted the laurdan emission in a dose-dependent manner (Fig. 5). Best fits to a single-site model indicate lower affinity (Kd=32 ± 4 μM, n=3) for isoform1 relative to the longer splice isoform 2 (Kd=14 ± 5 μM, n=3). We also measured a construct composed of a.a. 1-209 encoding both the C2A domain and adjacent linker and found that the inclusion of the domain did not shift the titration curve significantly (Fig. 5). This result agrees with our cosedimentation studies and suggest residues 1-140 including the C2A domain does not contribute to binding.

**Figure 5.**
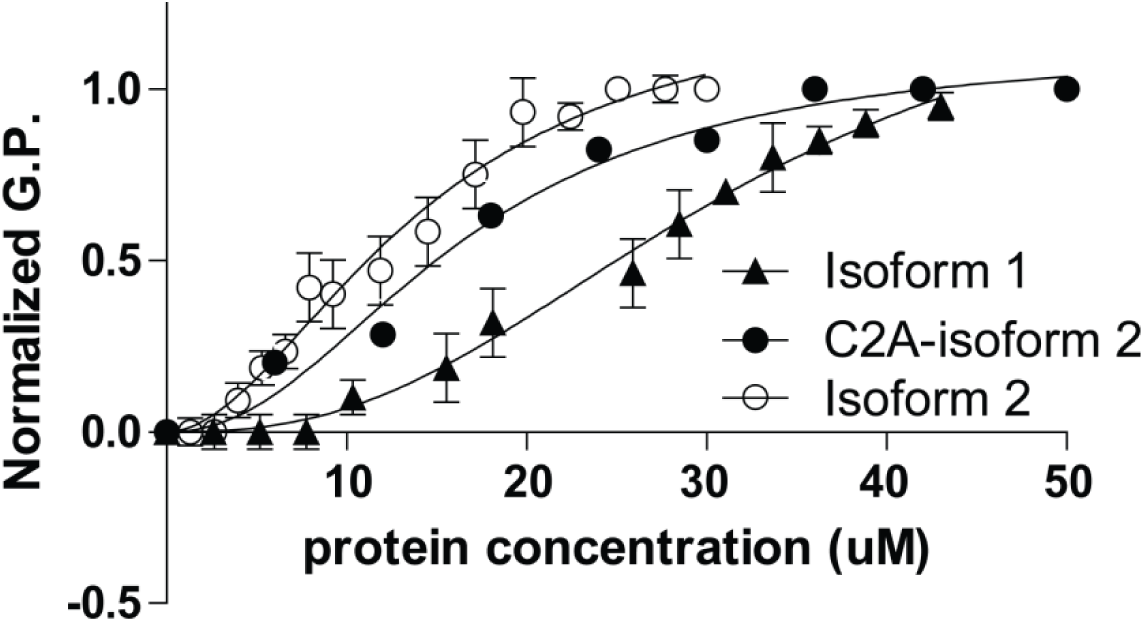
Both isoforms of the otoferlin C2A-C2B linker bind POPC/POPS liposomes. Titrations of isoform 1 linker, isoform 2 linker, and C2A-isoform 2 linker are plotted. Error bars represent S.D. n = 3.

We conclude that the C2A domain and N-terminal linker residues (a.a. 1-140) do not interact with membranes, while residues 141-209 of the C2A-C2B linker contains a sequence that directly interacts with negatively charged lipids.

### The linker is the major contributor to the membrane binding activity of the C2A- C2B region of otoferlin

Previous studies have established that the C2A domain of otoferlin does not bind membranes and the C2B domain binds with low affinity^31^. To determine the contribution of the linker to membrane binding when coupled to the flanking C2A and C2B domains we tested a construct composed of both C2 domains and linker. When tested we found that the construct bound liposomes in a Ca2+ insensitive manner with a dissociation constant (Kd= 18 ±1μM ± 4 μM, n=3) similar to the 141-209 linker construct (Fig. 6). By contrast C2A did not shift the laurdan fluorescence emission up to the limit of the titration, in agreement with results of our cosedimentation assay.^31^ Titration of the C2B domain yielded changes in the laurdan fluorescence at high concentrations of the domain in agreement with previous studies which reported a low affinity interaction.^31^ Thus the disordered interdomain linker is the major contributor to the membrane binding properties of the C2A-C2B N-terminus of otoferlin.

**Figure 6.**
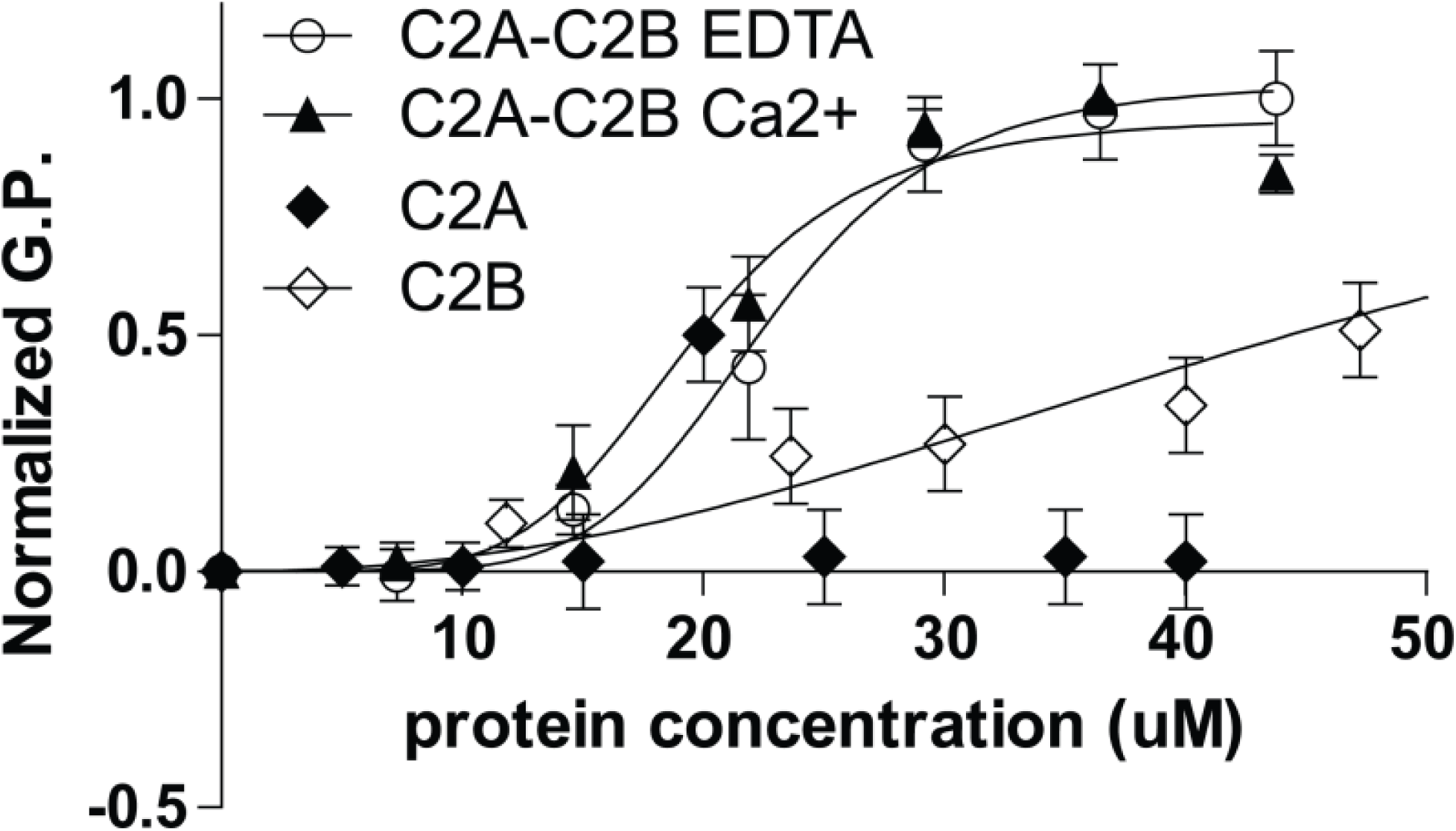
The otoferlin C2A-C2B linker contributes to the membrane binding activity of the N-terminal C2A-C2B region. Shown are titrations of single C2A domain which did not alter the fluorescence, as well as titrations of the C2B domain and C2A-C2B. Addition of otoferlin C2A-C2B shifts the G.P. value independent of Ca2+. *Error bars* represent S.D. *n* = 3.

### Electrostatics mediate C2A-C2B linker-membrane interaction

The otoferlin C2A-C2B linker sequence is enriched in charged and polar residues including serine, arginine, glutamic acid, and aspartic acid. This enrichment suggests an electrostatic basis for the preferential interaction with charged lipids. To assess the influence of electrostatics, we tested for linker-liposome interaction under different salt concentrations (Fig. 7). Compared to samples containing physiologically relevant NaCl concentrations (150 mM), measurements of samples containing 50 mM NaCl displayed greater binding sensitivity and an increase in the slope of the curve (Fig. 7). Measurements of samples containing 400 mM “high” salt conditions had the opposite effect, leading to less sensitivity and a gentler slope of the binding curve (Fig. 7). These results agree with an electrostatic basis for linker-membrane interaction.

**Figure 7.**
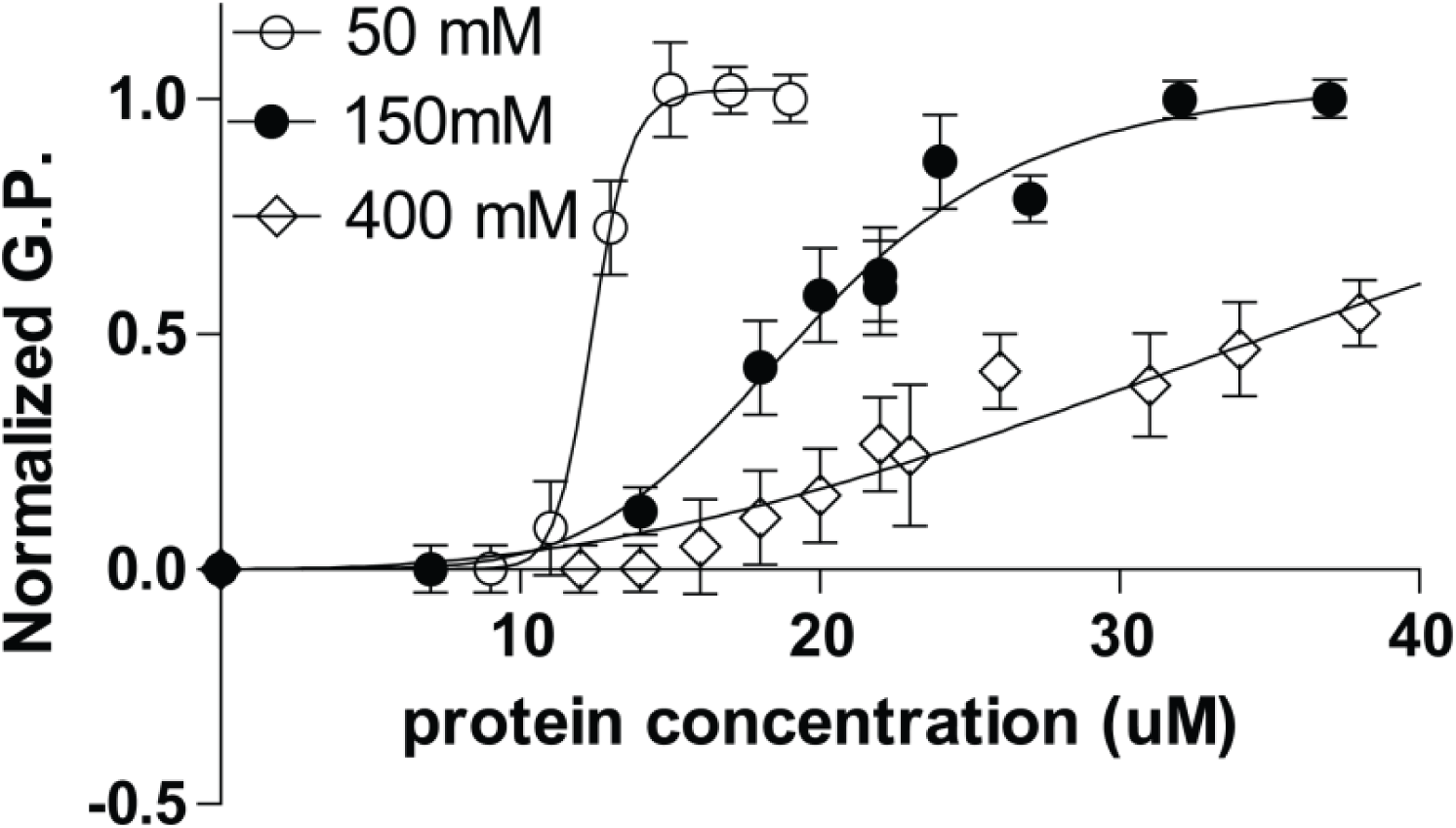
The interaction between the C2A-C2B linker and membranes is mediated by electrostatics. Titrations of the linker under 50 mM, 150 mM, or 400 mM NaCl are shown. Error bars represent S.D. n = 3.

To gain additional insight into the observed interaction we performed coarse-grained (CG) simulations of both otoferlin linker isoforms in the presence of a POPS/POPC lipid membrane (Fig. 8). For comparison we also included the otoferlin C2A domain and the dysferlin C2A-C2B linker. Relative to all-atom simulations, CG models offer a significant computational speedup due to the fewer number of interaction sites between molecules and faster configurational sampling due to the smoother free energy landscape.^15^ The latter can often result in sampling three to ten times faster compared to all-atom systems.^14,15^ The SPICA force-field has been shown to accurately reproduce structural, mechanical, and thermodynamic properties of lipid bilayers.^15^ It also reproduces dimerization free energies of transmembrane helices and peptides in solution, membrane-protein interactions, and radius of gyration for intrinsically disordered proteins.^14^

**Figure 8.**
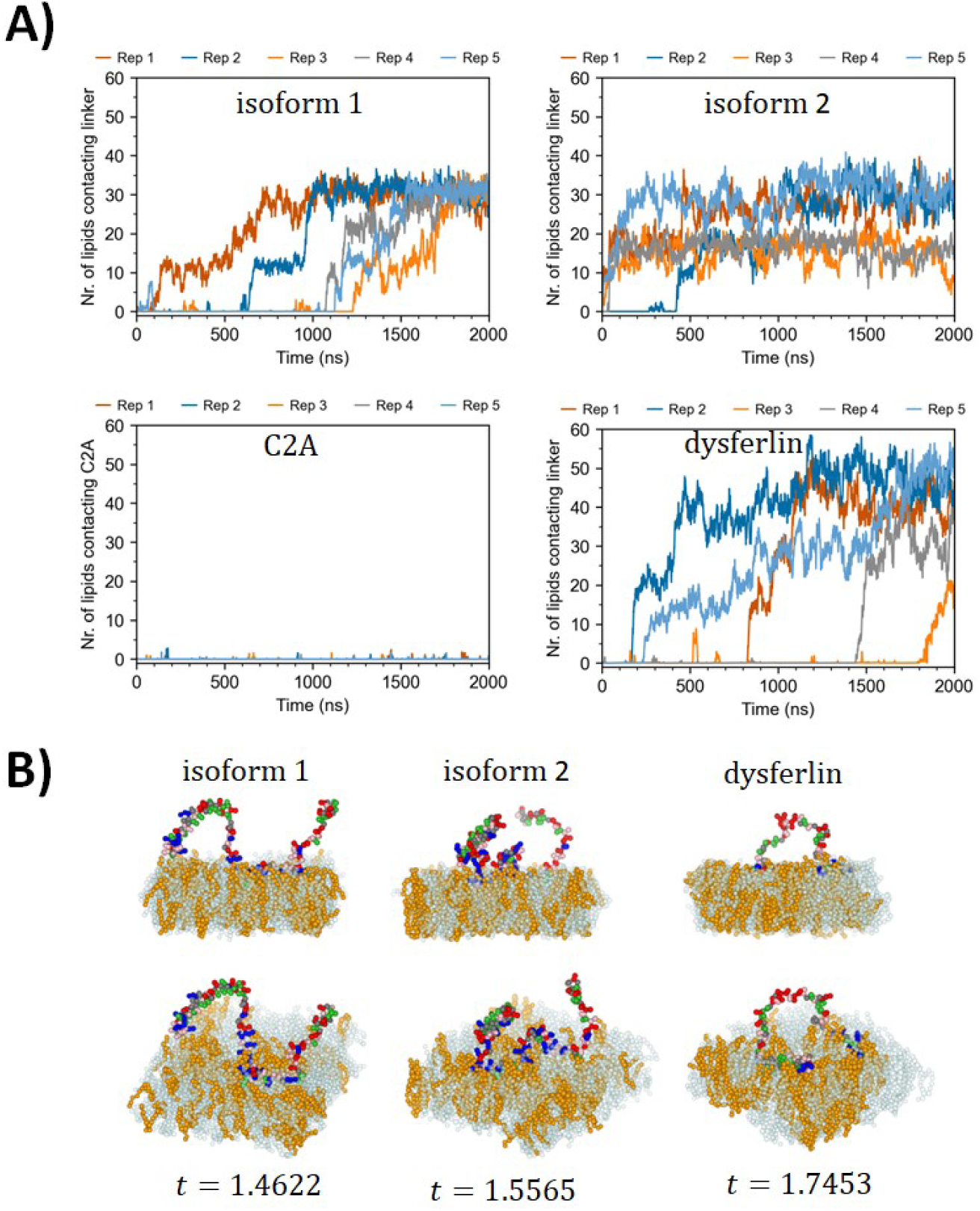
Coarse-grained MD simulations capture the binding interactions of otoferlin linker with lipid membranes. (A) Number of lipids contacting the linker for POPC:POPS membranes showing the increased ability of isoform 2 to interact with the membrane. Isoform 2 interacts with lipids faster and form a larger number of contacts. None of the 5 replicates for C2A displayed appreciable contacts, and dysferlin displayed heterogeneity in membrane association. (B) Side (top) and perspective (bottom) views of representative simulation trajectory frames of linkers interacting with the POPC/POPS membranes. Color legend: POPS:gold/orange, POPC:light blue/transparent, cationic residues (ARG, LYS, HIS):blue, anionic residues (GLU, ASP):red, polar (GLN, ASN, SER, THR):pink, hydrophobic (VAL, LEU, ILE, MET, PHE, TYR, TRP):green, other protein residues:gray. Regions with high frequency of anionic.

We first equilibrated the linker using an all-atom molecular dynamics simulation and subsequently applied the resulting energy-minimized structure in the SPICA coarse-grained simulation package. For all 5 replicates we found that over the span of the 2,000 nsec simulation time both otoferlin linkers increased the number of contacts with lipids in the membrane and associated with membranes using positively charged residues. However isoform 1 displayed slower kinetics and more instances of non-productive binding events, which we define as a transient contact with at least one lipid but less than ten that does not progress to full binding (Table 1). When the simulation was conducted with the C2A domain no significant binding was observed, in agreement with the cosedimentation and fluorescence results (Fig. 8). The dysferlin linker displayed considerable variability between the 5 replicates, however in all simulations the linker bound membranes with slower rates and a larger number of non-productive events than either otoferlin isoform 1 or 2 (Table 1). We conclude that the additional residues encoded by isoform 2 increase the number of successful contact events with the membrane resulting in higher rates of membrane binding.

**Table 1:** Number of non-productive binding events observed over the 5 simulation replicates.

|  |  |
| --- | --- |
| C2A | 64 |
| Isoform 1 | 43 |
| Isoform 2 | 1 |
| Dysferlin | 76 |

## Discussion

Ferlins are membrane trafficking proteins consisting of multiple Ca2+ sensitive domains interspersed between intrinsically disordered regions, including the linker between the C2A and C2B domains. Our comparison of vertebrate ferlin sequences suggests a conservation of the presence of this linker among family members, but with divergent sequence compositions that undergo alternative splicing. For otoferlin, alternative splicing of the C2A-C2B linker inserts additional residues that enhance the membrane binding activity of the linker region. Alternative splicing of the dysferlin also lengthens the linker, however the added sequence appears to introduce an acidic dileucine AP2 binding motif as identified using ELM. Splicing of the C2A-C2B linker of human myoferlin is also known to occur, and we therefore suggest that ferlins alter the C2A- C2B linker sequence to tune the number and type of binding motifs within the protein. Regardless of splicing, vertebrate ferlin C2A-C2B linkers are enriched in acidic and depleted in hydrophobic residues, and our measurements of otoferlin and dysferlin using CD and solution NMR are consistent with a disordered structure (Fig. 4).

We found the segment 140-209 of the otoferlin C2A-C2B linker bound membranes that included phosphatidylserine (Fig. 2,3). The observed interaction was found to be sensitive to ionic strength suggesting an electrostatic basis for binding (Fig. 6). Consistent with an electrostatic interaction, the otoferlin linker is hydrophilic with a calculated grand average hydropathy (GRAVY) of −1.0 for isoform 1 and −0.99 for isoform 2, and an enrichment of positively charged residues including 6 lysines and 7 arginines which in our simulations mediated the binding event (Fig. 7). Arginine-rich sequences are commonly found in many membrane binding and penetrating peptides where the charged guanidinium group interacts with charged components of the membrane. For comparison, the dysferlin linker has a calculated GRAVY of −0.68 suggesting a less hydrophilic (more hydrophobic) sequence relative to otoferlin.

In addition to lipid membranes, the otoferlin C2A-C2B linker is reported to bind AP2 via a dileucine motif located several residues N-terminal from the membrane interacting region (Fig. 2B, inset).^32^ Given the short distance between the dileucine motif and membrane binding region of the otoferlin linker we propose that otoferlin participates in vesicle recycling by placing the motif near the membrane for AP2 recruitment. Similarly, dileucine motifs are often found in the cytoplasmic tail of membrane proteins where the proximity of the motif to the membrane is proposed to promote AP2 membrane recruitment and activity. For example the cytoplasmic C-terminus of CD4 encodes a dileucine sequence proximal to the transmembrane domain which engages AP2 during endocytosis.^33^ Several spacer residues found between the binding motif and the end of the transmembrane domain of CD4 are required for optimal AP2 binding, and a similar spacing requirement may account for the residues between the AP2 and membrane binding regions of otoferlin C2A-C2B (grey linker residues between yellow and green segments in Fig. 2B inset). Since other ferlin C2A-C2B linkers also contain SLiMs and encode membrane binding segments we suggest that our proposed mechanism of endocytotic protein recruitment to the membrane may be a general function among family members.

