## Supplemental figures 1-4 for "Ferlin C2A-C2B linkers are alternatively spliced, intrinsically disordered, and interact with negatively charged membranes"

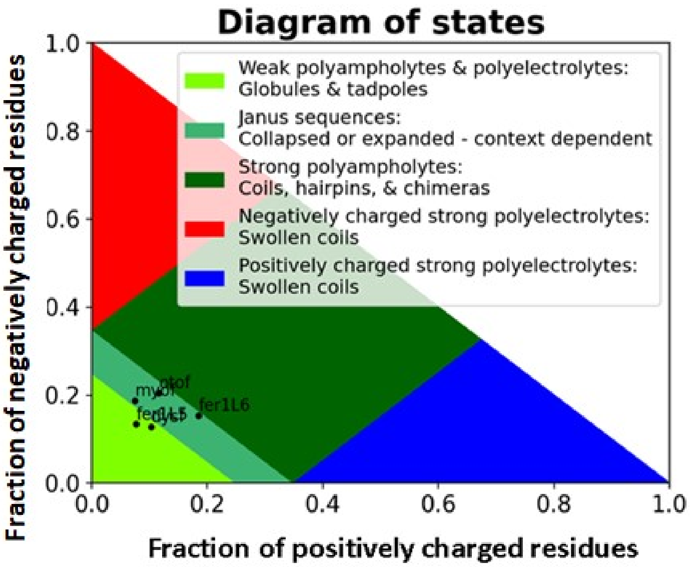


**Supplementary Figure 1:** The vertebrate ferlins plotted in a pappu diagram of states.

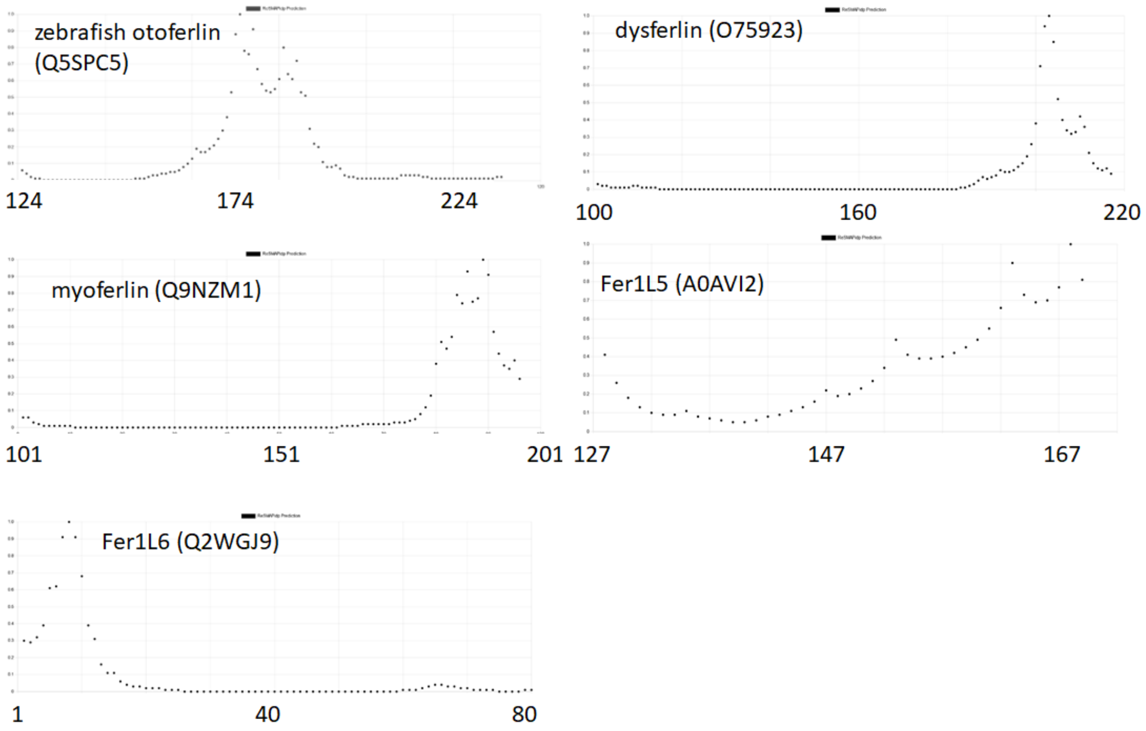


**Supplementary Figure 2**: ReSMAP plots of the C2A-C2B linker for zebrafish otoferlin, human dysferlin, myoferlin, Fer1L5 and Fer1L6.

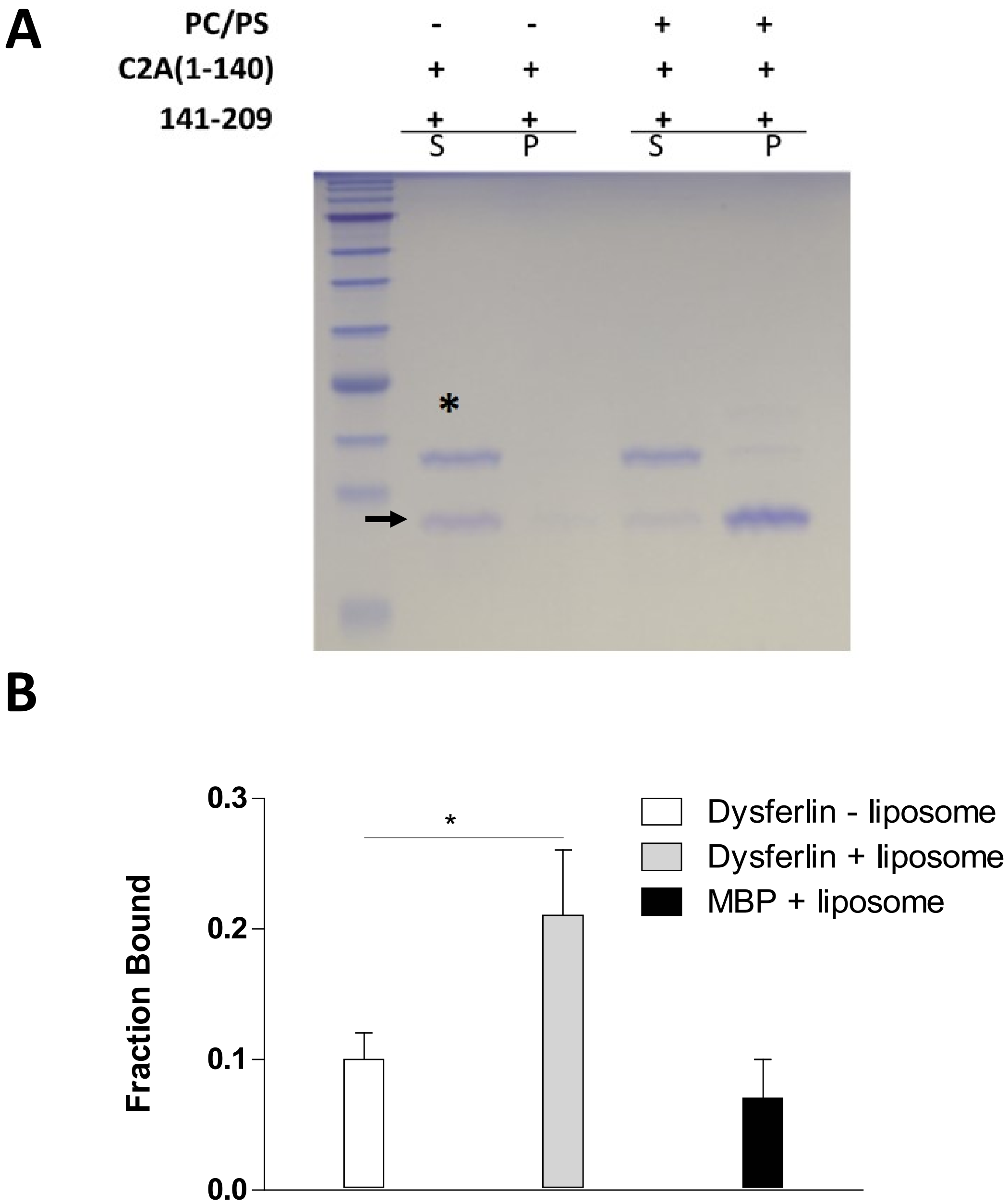


**Supplemental Figure 3**: (A) Otoferlin linker region 141-209 does not pull-down the N-terminus (amino acids 1-140) during cosedimentation with PS/PC liposomes. The first lane shows a molecular weight maker, followed by supernatant (S) and pellet (P) for samples. The band representing a.a. 1-140 (C2A) is denoted with an asterisk, and the membrane binding linker 141-209 denoted with an arrow. (B) Quantitation of the cosedimentation results of the liposome binding assay for the dysferlin linker with and without POPC/POPS liposomes. The maltose binding protein (MBP), which does not bind POPC/POPS liposomes, served as a control (t test, $p < 0.001$). Error bars represent ± standard deviation, $n = 3$.

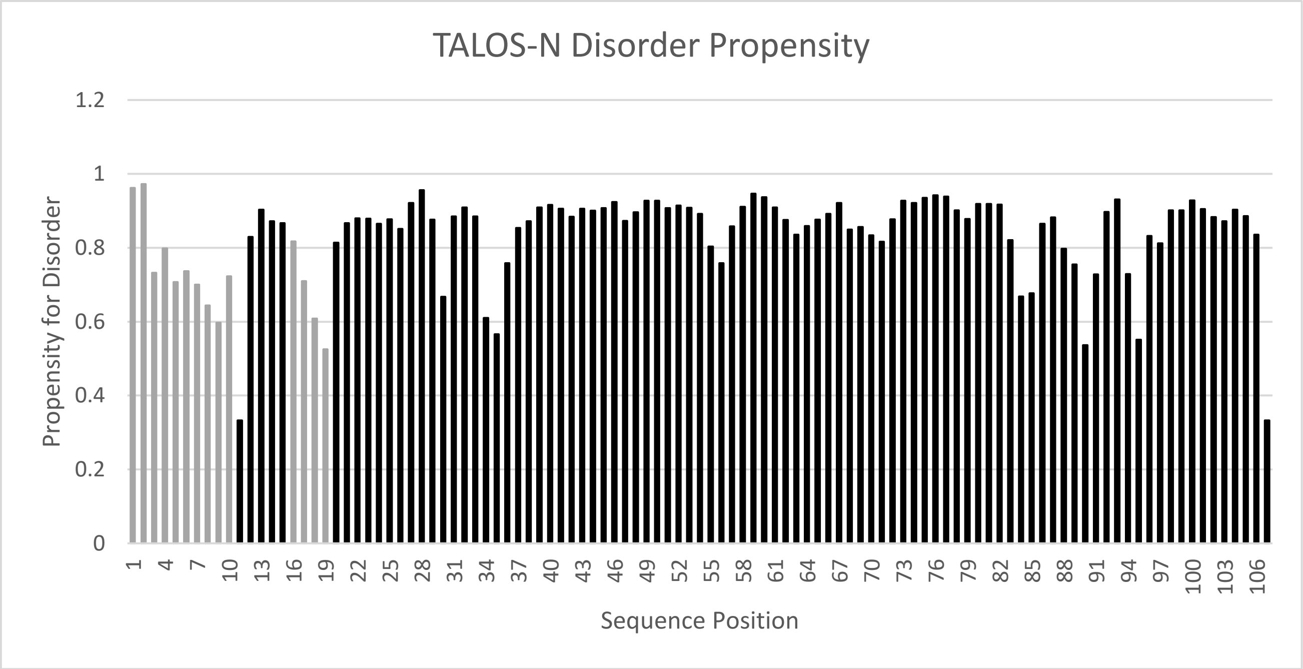


MGSSHHHHHHSSGLVPRGSHMASMTGGQQMGRGSEFEEKDSQETDGLLPGSRPSTRISGEKSFRSKGREK

TKGGRDGEHK AGRSVFSAMKLGKTRSHKEE PQRQDWW

**Supplemental Figure 4:** TALOS-N plot of the recombinant C2A-C2B linker fused to a uncleaved TEV cleavage sequence and polyhistidine purification tag at the N-terminus. Values determined using chemical shifts are denoted in black, and grey secondary structure propensity values for residues without assignment are based on the sequence alone.
